# Reinforcement learning in Parkinson’s disease is not associated with inflammatory tone

**DOI:** 10.1101/2023.09.28.557192

**Authors:** Jorryt G. Tichelaar, Marcel M. Verbeek, Iris Kersten, Roshan Cools, Rick C. Helmich

## Abstract

Parkinson’s disease (PD) is associated with large variability in the development and severity of both motor and nonmotor symptoms, including depression and impulse control disorder. Neuroinflammation might contribute to this heterogeneity, given its association with dopaminergic signalling, neuropsychiatric symptoms, and reward versus punishment learning. Here, we assessed the effect of inflammatory tone on probabilistic reinforcement learning and impulse control disorders in PD. We measured computational learning model-based neural reward prediction error and expected value signals in frontostriatal circuity during reinforcement learning using functional MRI. In addition, we acquired cerebral spinal fluid of 74 PD patients and screened for 13 inflammatory factors, including our primary marker of interest IL-6, previously implicated in reward learning signaling in the ventral striatum. In contrast to our prediction, we found no association between inflammatory tone and any of the behavioural or neural reinforcement learning parameters. Furthermore, we did we not find any correlation between inflammatory tone and depressive or impulsive PD symptoms. Exploratory analyses revealed a negative association between MCP-1 and reward prediction error signals in the ventral striatum, an observation that should be replicated in future work. The null findings might reflect the fact that measurements were taken ON medication, or that our sample consists of an early disease stage cohort that may be too small to detect these effects, or that IL-6 is a suboptimal marker for inflammatory tone, or a combination of these factors.

## Introduction

Parkinson’s disease (PD) is the fastest growing neurodegenerative disorder worldwide, characterized by severe dopamine depletion in the striatum.^1^ Patients suffer from the cardinal motor symptoms bradykinesia, rigidity and tremor, as well as non-motor symptoms such as autonomic dysfunction, neuropsychiatric symptoms and cognitive dysfunction.^2^ There are large inter-individual differences in the type and severity of symptoms that patients develop, and it is insufficiently clear where these differences originate from.^3,4^ In part, this heterogeneity is caused by inter-individual differences in how patients respond to dopaminergic medication.

Patients with PD are treated with levodopa and dopamine receptor agonists, which can alleviate motor and cognitive symptoms.^5^ Simultaneously, dopaminergic medication can also contribute to cognitive deficits, such as increased errors on associative learning tasks^6,7^ and reversal learning tasks.^8–11^ Such deficits are explained by pharmacological overstimulation of relatively intact dopaminergic reward pathways (see Cools et al.^4^ for a review), leading to an oversensitivity for rewards and reduced sensitivity to punishments.^12–14^ In severe cases, patients develop impulse control disorders (ICDs) that can manifest as pathological gambling, eating, shopping and/or hypersexuality.^15,16^ In terms of quality of life, ICDs are among the most impactful symptoms.^17–19^ Approximately 14% of PD patients develop ICDs, often after starting with or increasing the dose of dopamine receptor agonists.^15^ We have previously shown that PD patients with ICDs show a dopaminergic medication-related increase in choosing the rewarded versus punished option in a probabilistic reinforcement learning paradigm.^20,21^ This indicates the importance of altered dopaminergic reward signaling in ICDs. The obvious next question is what molecular mechanisms make some patients, but not others, vulnerable to medication-related overstimulation of the reward system?

There are three lines of evidence that suggest a key role for inflammatory tone in the pathophysiological mechanism of ICD in PD: 1) inflammation reduces dopamine synthesis capacity, 2) inflammation affects reward learning, and 3) patients with Parkinson’s Disease have a higher inflammatory tone.First, inflammation is associated with changes in dopamine availability, most likely because the dopamine synthesis pathway is particularly susceptible to reactive oxygen species, through oxidation of the cofactor BH4.^22^ In this way, inflammation may contribute to (further) dopamine depletion.^23–25^ This results in psychomotor slowing and cognitive deficits in many conditions associated with inflammation (see Felger et al.^26^ for a review). Second, inflammation affects the reward circuitry. Foremost, higher inflammation is associated with a blunted reward response in the ventral striatum.^27–29^ The ventral striatum is known to signal reward prediction errors (RPE),^30,31^ i.e. the difference between an expected and received reward. Indeed, Harrison et al.^32^ have shown reduced striatal reward prediction errors after a typhoid vaccination, compared with a placebo group.^32^ Third, increased inflammatory tone has been consistently found in PD. Compared with healthy controls, PD patients show increased activation of microglia,^33–36^ resulting in increased concentrations of pro-inflammatory cytokines^37–41^ and reactive oxygen species.^25,42,43^ Higher concentration of inflammatory cytokines are associated with increased neuropsychiatric suffering in PD,^44^ and have been hypothesized to actively contribute to (dopaminergic) cell death.^45,46^ Moreover, IL- 6, a key cytokine in the inflammatory cascade,^47,48^ has been found to be elevated in prodromal PD.^49^

Here, we tested whether inter-individual differences in inflammatory tone in PD patients are associated with abnormal reward versus punishment learning and clinical neuropsychiatric deficits, particularly ICDs. More specifically, we investigated: (1) associations between RL and inflammation in PD; (2) associations between ICDs and inflammation, and, in the presence of evidence for (1) en (2), we also assessed (3) whether any associations between ICD and RL are mediated by inflammation. We aimed to test the following three hypotheses. First, reward sensitivity, as measured by the response on an instrumental probabilistic RL task, is a function of inflammatory tone in PD patients. Second, increased impulsivity (higher QUIP-rs scores) and depression (higher BDI-II) are associated with higher inflammatory tone. Third, inflammation mediates the relationship between aberrant reinforcement learning and ICDs in PD. Additionally, since there is a vast body of literature indicating inflammation effects on depression,^27,28,50–52^ and since depression in PD is a risk factor for the development of ICDs,^53^ we also explored the effect of inflammation on depression in PD. We tested these hypotheses by acquiring cerebral spinal fluid in 74 PD patients from the Personalized Parkinson Project.^54^ This allows us to quantify inflammation directly at the site of PD neurodegeneration, instead of a indirect measure such as serum. Lerche *et al*. showed that associations between neurodegeneration and PD-related biomarkers were most robust in CSF compared to serum.^55^ Furthermore, they showed a sparse correlation between cytokine concentrations in the CSF and serum.^55^ Hence, we expected our approach to be more sensitive than previous studies.

We focussed on IL-6 as our primary cytokine of interest, because it is elevated in prodromal PD^49^ and a key player in the inflammatory cascade.^47,48^ Additionally, we explored the effects of Eotaxin, GRO-α, IL-8, IP-10, MCP-1, MIP-1β, RANTES, SDF-1α. These cytokines belong to the chemokine subgroup, which has previously been associated with neuropsychiatric symptoms.^44^ Lastly, we investigated the effect of IL-10, because of it’s anti- inflammatory property. All patients were clinically well-phenotyped in the neuropsychiatric, motor, and cognitive domains, and performed a probabilistic instrumental RL task in the fMRI scanner^21,54^. This allowed us to investigate whether naturally occurring variability in inflammatory markers in the CSF of PD patients contributes to inter-individual variability in value-based learning or choice, as well as neuropsychiatric symptoms.

## Materials and methods

This study is an extension of earlier work. Therefore, the general procedure, the instrumental learning task and MRI analysis are summarized, with more details to be found in Tichelaar *et al.*.^21^

### Ethics

This study was approved by the local institutional review board (Commissie Mensgebonden Onderzoek Region Arnhem-Nijmegen; reference number 2016–2934; NL59694.091.17) and was conducted in accordance with the Ethical Principles for Medical Research Involving Human Subjects, as defined in the Declaration of Helsinki (version amended in October 2013). All participants gave written informed consent.

### Participants

All data has been acquired through the Personalized Parkinson Project (PPP), which is a longitudinal observational study in 520 PD patients.^54^ Participants were included only when they have a diagnosis of idiopathic PD, were older then 18, <5 years have passed since diagnosis, and are without contraindications for MRI. Of the 520 patients, 141 completed the RL task analyzed in the current work. Of those, 74 patients underwent a lumbar puncture. To avoid effects of medication, we excluded three participants that did not yet use dopaminergic medication as treatment. Additionally, we excluded five participants who were unable to finish the RL task and four participants who did not complete the clinometric questionnaire battery, from the analysis in which those were required. Participants were reimbursed for a hotel stay on the night between the measurement days as well as their travel expenses. Furthermore, participants received 5% (€4.52 on average) of their total winnings from the RL task.

### General procedure

All PPP subjects underwent two measurement sessions across two days. Day 1 of the PPP consisted of the MRI session, including both the RL-task during fMRI scanning as well as a T1 anatomical scan. This session was ON their normal medication regime. Patients were offered an overnight stay in a local hotel, prior to the second testing day. Day 2 involved clinical assessment of their motor symptoms, while OFF their medication (i.e. after a 12h withdrawal prior to the onset of the measurements). After these initial assessments, patients resumed their standard medication regime and were tested ON medication, including both the motor test battery (i.e. UPDRS, including Hoehn&Yahr) and cognitive test battery (i.e. 15 words test, Benton Judgement of Line Orientation, Brixton Executive Functioning test, Numbers and Letters pronunciation test, Montreal Cognitive Assessment (MOCA), Semantic fluency & the Symbol Digit modalities test (SDMT), in addition to demographic and medical information). At home, participants were asked to complete a neuropsychiatric test battery, including the Beck Depression Inventory II (BDI), the Questionnaire for Impulsive-Compulsive Disorders in Parkinson’s Disease-Rating Scale (QUIP-rs) and the State Trait Anxiety Inventory for Adults (STAI). For a detailed description see Bloem et al.^54^

### Instrumental probabilistic learning task

To assess reward and punishment learning, all participants engaged in an instrumental probabilistic learning task. The task was based on Pessiglioni *et al.*,^31^ and adapted by Tichelaar *et al.*^21^ In short, participants were asked to maximize monetary output, by choosing between two cues with different monetary outcomes. There were two types of cue pairs: GAIN cues and LOSS cues. Participants were instructed to maximize reward. They did so by learning which cue was optimal and associated with reward (i.e. €10 in GAIN trials, €0 in LOSS trials) in 75% of the trials and with punishment (i.e. €0 in GAIN trials, -€10 in LOSS trials) in 25% of the trials. Simultaneously, participants had to avoid the suboptimal cue, which had the reversed reward to punishment outcome probability (i.e. 25%:75% trials). Different cues were used for GAIN and LOSS trials. Participants took one training run outside the scanner, and two runs within the scanner. Each run contained 28 GAIN trials, interleaved with 28 LOSS trials, see Fig 1.

**Figure 1.**
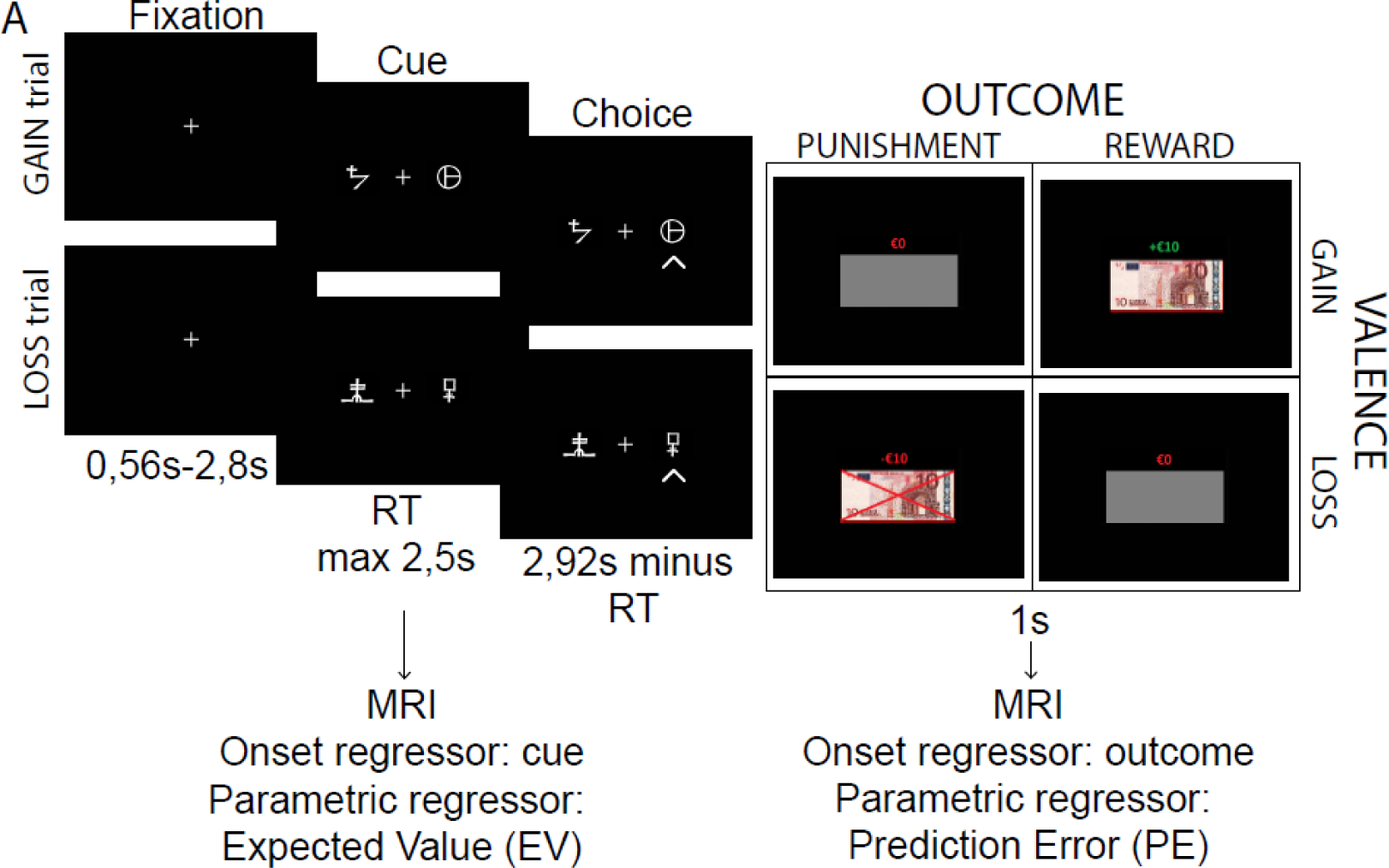
Task description and a Win-Stay-Lose-Shift trial explanation – (A) Participants performed a probabilistic instrumental learning task according to Pessiglione et al..

### CSF sample collections

A subset of the PPP cohort voluntarily agreed to a lumbar puncture. Lumbar punctures were performed under local anesthesia and with the standardized protocol of the Radboudumc. Samples were frozen at -80 degrees Celsius in the local Biobank.

Upon retrieval of the samples from the Biobank, we used the Chemokine 9-Plex Human ProcartaPlex™ Panel 1 (Catalog number: EPX090-12187-901) to determine the concentration of Eotaxin (CCL11), GRO-α (CXCL1), IL-8 (CXCL8), IP-10 (CXCL10), MCP-1 (CCL2), MIP-1α (CCL3), MIP-1β (CCL4), SDF-1α and RANTES (CCL5). Furthermore, the High Sensitivity 9-Plex Human ProcartaPlex™ Panel (Catalog number: EPXS090-12199-901) was used to determine the concentration of IFN gamma, IL-1β, IL-2, IL-4, IL-6, IL-10, IL-12p70, IL-17A (CTLA-8) and TNF-α. Both kits were run in duplicate per participant and in accordance with the manufacturer’s manual. One exception was made for the chemokine panel. Instead of a 60 to 120-minute antigen incubation at room temperature, the antigen incubation was done overnight at 4 degrees. This was recommended for samples which require high sensitivity.

Fluorescent intensity was measured by the Luminex MagPix system met xPONENT software (xPonent build 4.2.1324.0, Luminex cooperation, Austin, TX, USA) and analyzed with Millipore Milliplex Analyst version 3.5.5.0. Concentrations were determined with the use of baseline fluorescent curves as calculated with known concentration samples as provided by the manufacturer.

### CSF marker exclusion

After analysis of the CSF samples, we excluded MIP-1α, IL-2 & IL-12p70 from analysis, because our quality control samples (in-house samples, used to compare between kits) and all subject samples were below the lowest limit of quantification. Furthermore, all samples for IL- 1β, IL-4, IL-17A, IFN-γ & TNF-α were below the absolute detection threshold and returned blank. Thus, our current experimental setup was insufficient to reliably quantify MIP-1α, IL- 2, IL-12p70, IL-1β, IL-4, IL-17A, IFN-γ & TNF-α.

All samples were measured in duplicate. Ten samples differed more than 20% between the two duplicate measurements, but were still included in the analysis. Similarly, GRO-α, MIP-1β, RANTES, IL-6 and IL-10 contained samples in which the concentration was below the lowest limit of quantification, but above the absolute detection threshold, were also still included in the analysis.

### Statistical analysis

#### Correlations

We assessed frequentist correlation between the clinometric scores, mean BOLD response and the cytokines by calculating the spearman rho partial correlation (“partialcorr” function, MATLAB^56^ version 2022b). Partial correlation was used to correct for confounding effects of age, sex and BMI. Additionally, we performed a false discovery rate analysis, to correct for multiple comparisons, according to the Benjamini–Hochberg procedure with a false discovery rate of 20%. To assess the evidence for null-results, we additionally assessed a Bayesian variant of the spearman rho, according to van Doorn et al.^57^ We used R version 3.6.1^58^ and the software provided by van Doorn et al.,^57^ available at https://osf.io/gny35/.

#### Reinforcement learning behavior

We assessed three behavioral readouts of the RL-task. First, we assessed accuracy, defined as the percentage of trials on which the most rewarding cue was chosen. Second, we assessed win-stay-lose-shift behavior, defined as the percentage of trials in which the participant chose the same cue after a reward, or switched cue after a punishment. Third, we assessed reaction time.

RL parameters were investigated using Bayesian mixed effects logistic regression on per trial accuracy, stay behavior and reaction time. To do so, we used R version 3.6.1^58^ and the brm function from the brms package.^59^ The model space is presented in supplementary table 2 and the regressors included in our models are presented in supplementary table 3. All continuous variables were z-scored to increased interpretability of the beta values. All variables were added as fixed effects and additionally valence and outcome were added as random slopes for each subject, because their values change over trials. We used the default brms priors and the models were fit using four chains with 6000 iterations and 1500 warmup iterations. Accuracy and WSLS behavior were assessed using the Bernoulli family with a logit link. Trial wise reaction times were analyzed with a Gaussian family, with identity link.

### MRI analysis

Preprocessing and first level analysis was performed identical to Tichelaar *et al.*^21^ In short, BOLD images were acquired with a five-echo sequence with a TR of 2240ms. The images were preprocessed using fMRIprep 20.2.1.^60^ and smoothed with a 8mm kernel. Based on the behavioral choices during the reinforcement learning task, we calculated a per trial RPE and expected value (EV) regressor, according to a standard Rescorla Wagner model:

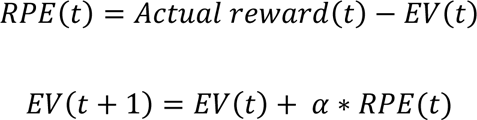

Where t is the current trial, RPE is the reward prediction error, actual reward is the reward outcome (i.e. €10, €0 or -€10), EV is the expected value and alfa is the learning rate (fixed to 0.2 to avoid MRI differences based differences in learning rate^61^). We added the RPE regressor as parametric modulator to the onset regressor for outcome. The EV regressor was added to the onset regressor at the time of choice. For both outcome and choice, we had two onset regressors, one for GAIN trials and one for LOSS trials. Lastly, we added 46 motion regressors and the AROMA regressors that did not correlate with the task regressors, as well as two intercept regressors denoting the two task blocks.

Our primary analysis consisted of the ranked correlation between natural log of the cytokine concentrations and the mean first level beta values for two, a priori chosen region of interest (ROI). The ROI was based on a meta-analysis of RPE and EV activation during reinforcement learning paradigm, see supplementary material.^62^ The RPE ROI consisted of the striatum, left insula and part of the thalamus, V3 and V4. The EV ROI consisted of the vmPFC. Through SPM, we retrieved the beta values of both the RPE and EV ROI’s for the GAIN trials only, LOSS trials only and GAIN > LOSS contrast. The beta values were averaged across the ROI and correlated with the natural log of the concentration of the cytokines.

To further assess whether group level activation patterns were a function of MCP-1 or IL-6, we ran a second-level random effects analysis. MCP-1 was chosen because of the correlation with the RPE BOLD signal, and IL-6 was chosen because we were primarily interested in this cytokine (see introduction). We used a linear regression analysis over all subjects, with the natural log of the IL-6 or MCP-1 concentration as covariate. Multiple comparison correction was performed through threshold-free cluster enhancement (TFCE).^63^ We used 5,000 permutations and reported significant clusters with a *P*_fwe_<0.05, unless stated otherwise.

### Data availability

The final codebase used for this paper is available at the Donders Repository: https://doi.org/10.34973/13w4-gw70. The research data is available on request only, to ensure the privacy of the participants. A data acquisition request can be sent to.

## Results

### Demographics

Mean demographic information and cytokine concentration across all subjects is displayed in Table 1.

**Table 1.** Patient Characteristics.

|  |  | RANGE | MEAN | STD |
| --- | --- | --- | --- | --- |
| DEMOGRAPHICS | Number of Participants |  | 71 |  |
|  | Age |  | 60.24 | 8.31 |
|  | Gender |  | 36 women,<br>35 men |  |
| MEDICATION | LEDD |  | 541.5 | 338.06 |
|  | Agonist Use |  | No:47,<br>Yes:24 |  |
| PD SEVERITY | UPDRS-III (OFF med) | 0-123 | 30.68 | 11.29 |
|  | Months Since Diagnosis |  | 26.03 | 18.08 |
|  | Dyskinesias |  | No:70, Yes:1 |  |
| NEUROPSYCHIATRY | Depression (BDI-II) | 0-63 | 9.39 | 7.95 |
|  | Anxiety (STAI) | 0-80 | 73.46 | 21.05 |
|  | Impulse Control Disorders (QUIP-rs) | 0-112 | 8.22 | 11.56 |
|  | Cognitive Functioning (MOCA) | 0-30 | 28.32 | 2.03 |
| EXECUTIVE FUNCTIONING | Benton Task |  | 25.96 | 3.78 |
|  | Brixton Task |  | 12.83 | 6.21 |
|  | Numbers And Letters Task |  | 19.85 | 2.94 |
|  | Semantic Fluency |  | 26.24 | 4.84 |
|  | Symbols & Digits Task |  | 38.35 | 7.77 |
| INFLAMMATION | Eotaxin (pg/ml) |  | 0.83 | 0.13 |
| | GROα (pg/ml) | | 2.07 | 0.42 |
|  | IL-8 (pg/ml) |  | 9.76 | 4.9 |
|  | IP-10 (pg/ml) |  | 26.5 | 16.41 |
|  | MCP-1 (pg/ml) |  | 408.48 | 192.35 |
| | MIP-1β (pg/ml) | | 4.54 | 0.93 |
|  | RANTES (pg/ml) |  | 0.52 | 0.11 |
| | SDF-1α (pg/ml) | | 690.97 | 219.1 |
|  | IL-6 (pg/ml) |  | 2.76 | 2.45 |
|  | IL-10 (pg/ml) |  | 0.31 | 0.17 |

### Correlations between cytokines

We obtained the CSF concentration of ten cytokines, consisting of IL-6, IL-10, and eight chemokines (i.e. eotaxin, GRO-α, IL-8, IP-10, MCP-1, MIP-1β, RANTES and SDF-1α). We found significant correlations between the CSF markers, as shown in table 2. Foremost, the chemokines correlated with each other. This led us to calculate the first principal component (via PCA) over all chemokines, to capture the specific effect of chemokines in one variable. As expected, the first component correlated with all chemokines, but not with IL-6 (Rho(63)=0.07, p=0.56) or IL-10 (Rho(64)=0.21, p=0.20).

**Table 2.**
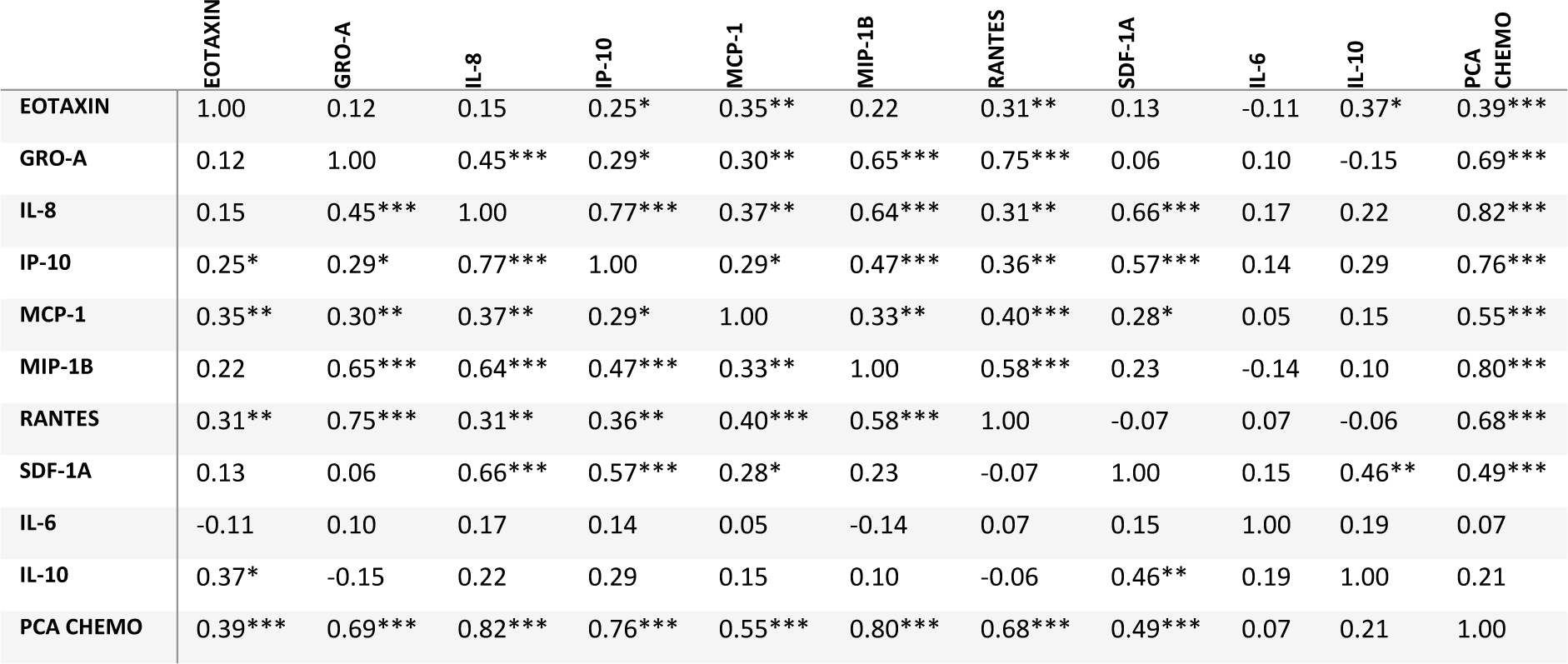
Correlations between cytokines. All Spearman Rho correlation between cytokine concentrations. *p<0.05, **p<0.01, ***p<0.001

|  | EOTAXIN | GRO-A | IL-8 | IP-10 | MCP-1 | MIP-1B | RANTES | SDF-1A | IL-6 | IL-10 | PCA CHEMO |
| --- | --- | --- | --- | --- | --- | --- | --- | --- | --- | --- | --- |
| EOTAXIN | 1.00 | 0.12 | 0.15 | 0.25* | 0.35** | 0.22 | 0.31** | 0.13 | -0.11 | 0.37* | 0.39*** |
| GRO-A | 0.12 | 1.00 | 0.45*** | 0.29* | 0.30** | 0.65*** | 0.75*** | 0.06 | 0.10 | -0.15 | 0.69*** |
| IL-8 | 0.15 | 0.45*** | 1.00 | 0.77*** | 0.37** | 0.64*** | 0.31** | 0.66*** | 0.17 | 0.22 | 0.82*** |
| IP-10 | 0.25* | 0.29* | 0.77*** | 1.00 | 0.29* | 0.47*** | 0.36** | 0.57*** | 0.14 | 0.29 | 0.76*** |
| MCP-1 | 0.35** | 0.30** | 0.37** | 0.29* | 1.00 | 0.33** | 0.40*** | 0.28* | 0.05 | 0.15 | 0.55*** |
| MIP-1B | 0.22 | 0.65*** | 0.64*** | 0.47*** | 0.33** | 1.00 | 0.58*** | 0.23 | -0.14 | 0.10 | 0.80*** |
| RANTES | 0.31** | 0.75*** | 0.31** | 0.36** | 0.40*** | 0.58*** | 1.00 | -0.07 | 0.07 | -0.06 | 0.68*** |
| SDF-1A | 0.13 | 0.06 | 0.66*** | 0.57*** | 0.28* | 0.23 | -0.07 | 1.00 | 0.15 | 0.46** | 0.49*** |
| IL-6 | -0.11 | 0.10 | 0.17 | 0.14 | 0.05 | -0.14 | 0.07 | 0.15 | 1.00 | 0.19 | 0.07 |
| IL-10 | 0.37* | -0.15 | 0.22 | 0.29 | 0.15 | 0.10 | -0.06 | 0.46** | 0.19 | 1.00 | 0.21 |
| PCA CHEMO | 0.39*** | 0.69*** | 0.82*** | 0.76*** | 0.55*** | 0.80*** | 0.68*** | 0.49*** | 0.07 | 0.21 | 1.00 |

We found no correlations between IL-6 and any of the chemokines, IL-10 (Rho(69)=0.19, p=0.23), or the PCA over all chemokines (Rho(69)=0.07, p=0.54). Furthermore, IL-10 showed a positive correlation with eotaxin (Rho(70)=0.37, p=0.02) and SDF-1α (Rho(70)=0.46, p=0.003), which is surprising due to the anti-inflammatory properties of IL-10.^64^

### No evidence for a link between IL-6 concentration and probabilistic instrumental learning in PD

Across participants we show successful engagement in the reinforcement learning paradigm, see supplementary material & Fig. 2A. Here we assessed whether RL behavior depended on IL-6. First, we assessed whether IL-6 concentration affected accuracy, i.e. the percentage of trials on which the most rewarding option was chosen. We found no effect on accuracy across all trials (correct response ∼ * ln(IL-6); brms 95% CI=[-0.14 0.32]). We also did not see an effect of IL6 on accuracy as a function of outcome valence (GAIN versus LOSS; correct response ∼ valence * ln(IL-6); brms 95% CI=[-0.29 0.10], Fig. 2B). Second, we assessed whether win-stay-lose-shift behavior changed as a function of IL-6 concentration. Win-stay-lose-shift behavior is defined as the percentage of trials in which the participants switch cue, after receiving a punishment, or stay with the cue after receiving a reward. Again, WSLS- behavior did not chance as a function of IL-6 across trials (Stay response ∼ outcome * ln(IL- 6); brms 95% CI = [-0.12 0.09]), or as a function of valence (Stay response ∼ valence * outcome * ln(IL-6); brms 95% CI = [-0.14 0.11], Fig. 2C). Third, there were also no effects of IL-6 on reaction times (response time ∼ ln(IL-6); brms 95% CI=[-0.03 0.01], Fig. 2D), also not as a function of valence; GAIN vs LOSS (response time ∼ valence * ln(IL-6); brms 95% CI=[-0.07 0.04]). Additionally, we explored the effects of all other cytokines and chemokines on either accuracy, RT or WSLS-behavior, but found no significant effects, see supplementary table 1.

**Figure 2.**
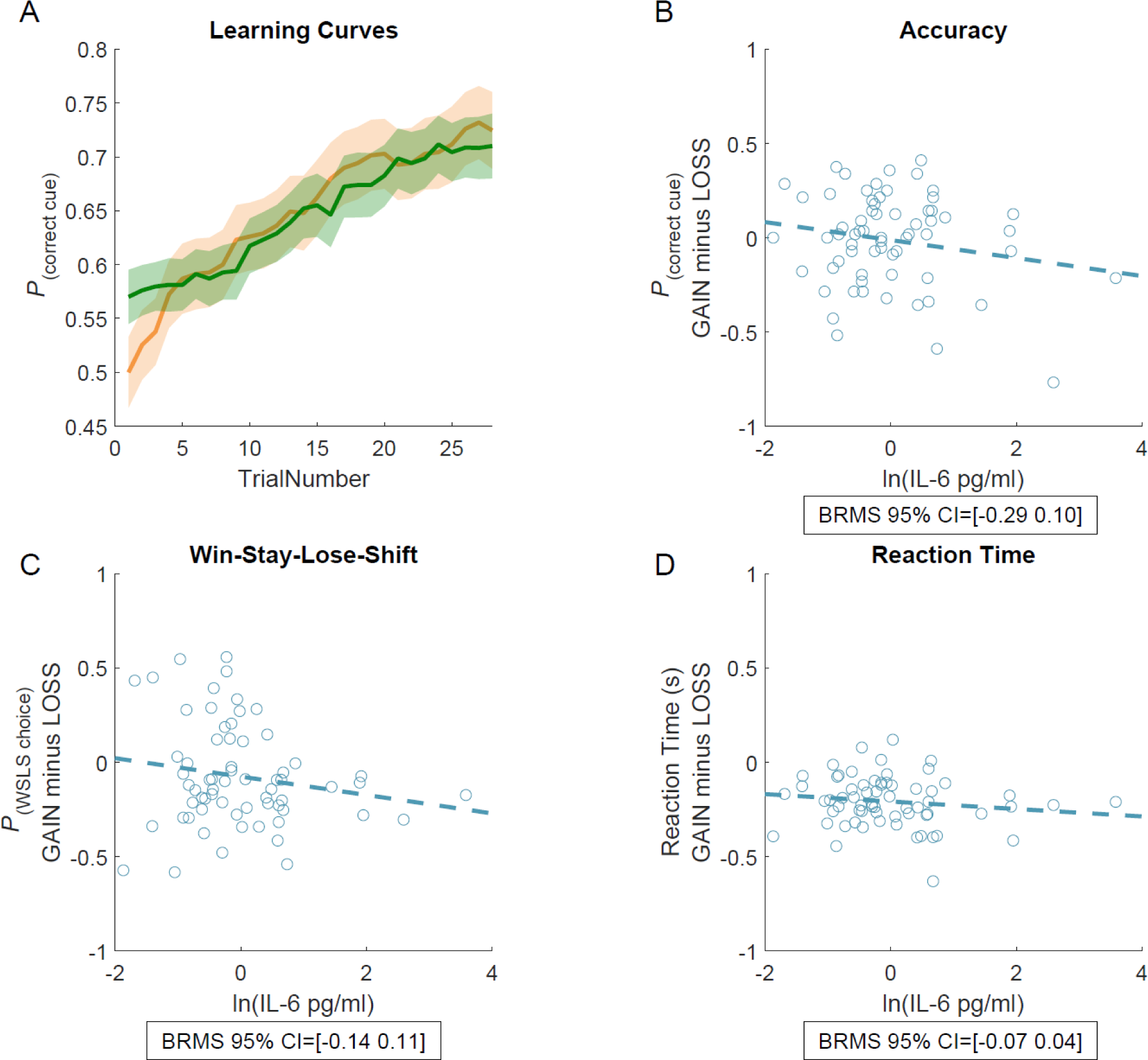
Reinforcement learning behavior. – **(A)** Across participants, accuracy increases over time for both GAIN (orange) and LOSS (green) trials. **(B)** Accuracy in GAIN trials, minus accuracy in LOSS trials as a function of the natural log of IL-6 concentration. **(C)** Win-Stay-Lose-Shift behavior in GAIN trials, minus LOSS trials as a function of the natural log of IL-6 concentration. **(D)** Reaction time in GAIN trials, minus LOSS trials as a function of the natural log of IL-6 concentration.

### IL-6 concentration is not associated with impulsivity or depression in patients with PD

In contrast to our hypothesis, we found moderate evidence *against* a correlation between IL-6 and the QUIP-rs score for ICDs (Bayesian Spearman Rho=-0.34, BF01=6.43, Fig. 3B). We also explored whether any of the other cytokines correlated with the QUIP-rs, but found not significant correlations (see supplementary table 1). Additionally, we assessed the correlation between depression (BDI-II score) and IL-6. Again, we found moderate evidence *against* such a correlation (Bayesian Spearman Rho=0.00, BF01=6.97, Fig. 3A). We also explored whether any of the other cytokines correlated with the QUIP-rs or BDI-II. We found a negative correlation between depression (BDI-II) and RANTES (Rho(70)=-0.27, p=0.03, Fig. 3C), but this did not survive correction for multiple comparison (FDR, Benjamini-Hochberg procedure, Q=0.2). In summary, we found evidence against a link between IL-6 concentrations and impulsivity or depression. For additional explorative correlations between any cytokine and any clinometric factor, see supplementary table 1.

**Figure 3.**
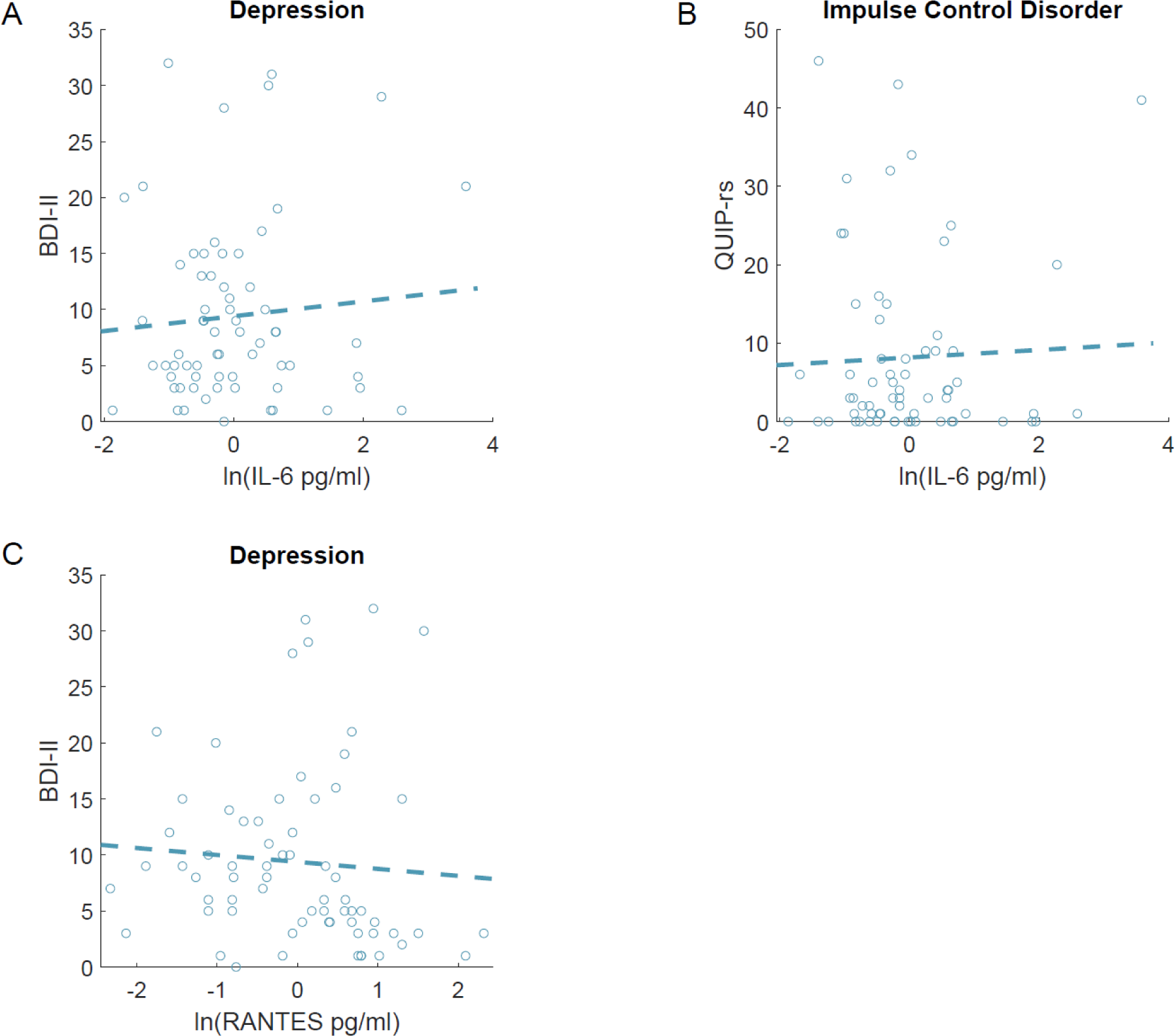
Correlations between cytokines and neuropsychiatric scores. – Here we depict the correlations between **(A)** depression (BDI-II) and IL-6 **(B)** impulsivity (QUIP-rs) and IL-6 **(C)** depression (BDI-II) and RANTES. The correlation between IL- 6 and BDI-II and QUIP-rs are not significant. The correlation between depression (BDI-II) and RANTES is (Rho(70)=-0.27, p=0.03), but does not survive multiple comparison correction and is not substantiated by a Bayesian correlation (BF01 = 0.58).

### No evidence for associations between IL-6 and neural reward prediction error or expected value signaling

We investigated the effect of inflammatory cytokines on RL related BOLD signal. In line with previous work,^62^ we found robust and significant activation of the reward network during the task. This includes a significant main effect of RPE in the ventral striatum and a main effect of EV in a ROI of the vmPFC, across all participants, see supplementary material.

To assess whether RPE/EV BOLD signaling is a function of IL-6, we correlated the first level beta values from the ROIs with IL-6 concentration. We found no correlation between IL-6 concentration and the beta values of the RPE (GAIN + LOSS; Rho(64)=-0.20, P=0.10) or EV (GAIN + LOSS; Rho(64)=-0.03, P=0.79), nor as a function of valence for RPE (GAIN > LOSS; Rho(64)=0.15, p=0.24, Fig. 4A) or EV (GAIN > LOSS; Rho(64)=0.03, p=0.80, Fig. 4B). Additionally, when adding IL-6 concentrations as a cofactor to our second level GLM, we did not find any activity that survived correction for family wise error correction.

**Figure 4.**
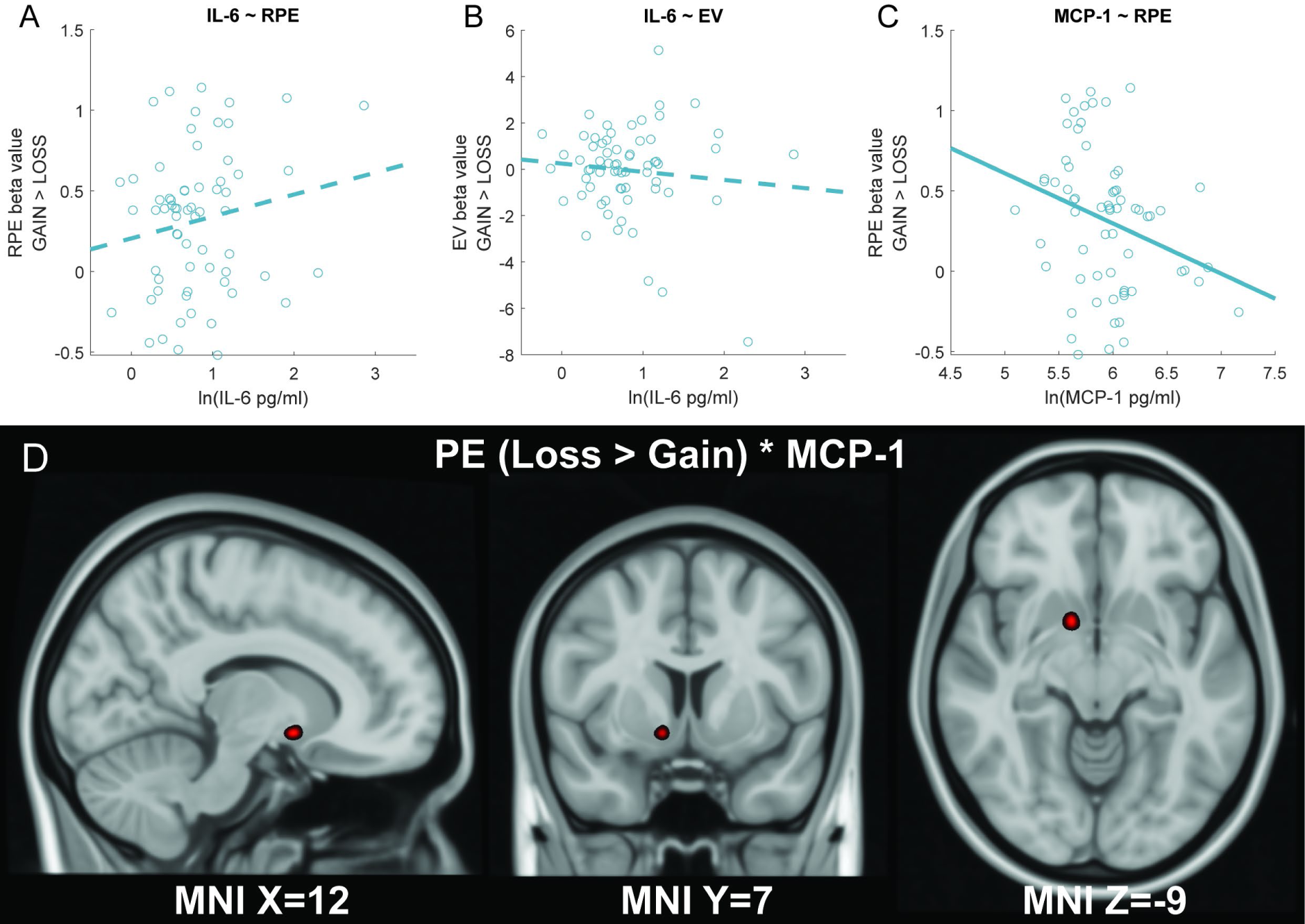
Differences in BOLD response as a function of IL-6 and MCP-1. – Correlation between the natural log of IL-6 concentration and **(A)** Mean beta values for prediction error (not significant) **(B)** Mean beta values for expected value (not significant). **(C)** Correlation between natural log of MCP-1 and mean beta values for prediction error (significant). **(D)** Reduced activation in the striatum for GAIN trials, compared to LOSS trials (i.e. LOSS > GAIN). Prediction error was added as parametric modulator on the first level, MCP-1 concentration as cofactor on the second level. TFCE fwe p<0.05.

Interestingly, when we explored the effects of other cytokines on reward prediction error signaling, we found that patients with higher MCP1 concentrations showed a significantly reduced GAIN > LOSS reward prediction error signals (Pearson Rho = -0.32, p=0.009, see Fig. 4C). However, the Bayes factor is near 1, indicating insufficient power (BF10=0.96) and the correlation does not survive multiple comparison correction. Second-level group analysis did reveal a small, but significant cluster in the ventral striatum as a function of MCP-1 (7 voxels, MNI local maximum [12 7 -9], TFCE = 51.43, P_fwe_ = 0.024, Fig. 4D). This suggests decreased RPE BOLD signalling during GAIN trials, compared to RPE signalling in LOSS trials, when patients have higher concentrations of MCP-1. We assessed whether this effect was associated with more depressive symptoms. However, we found no evidence for an association between MCP-1 concentration and depression (Perason Rho(65)=-0.17, p=0.17), nor between the beta values for the LOSS > GAIN association with MCP-1 and depression (Pearson Rho(60)=0.24, p=0.06).

## Discussion

The present work aimed to demonstrate that naturally occurring variance in cytokine concentrations is associated with ICD symptoms in PD patients ON medication, as well as with associated cerebral or behavioural indices of reinforcement learning . However, there was no evidence for such associations. Furthermore, there was no evidence that IL-6 is associated with symptoms of depression. When we explored correlations with MCP-1, we found that higher concentrations correlated with reduced neural reward prediction error signal during GAIN trials compared with LOSS trials. This effect should be interpreted with caution, because it did not survive correction for multiple comparisons. Future research is required to replicate the link between neural reinforcement learning signals and MCP-1.

To our surprise, we did not replicate previous findings that inflammation was associated with reinforcement learning, as previously shown in healthy controls.^32^ Furthermore, we find no correlation between inflammatory tone in CSF and neuropsychiatric symptom severity in PD (ICD or depression).^44^ This suggests that baseline variance in inflammatory tone in PD might not be sufficient to change the reward sensitivity, at least, not to the extent that it can be associated with changes on the reinforcement learning paradigm or lead to reward related neuropsychiatric conditions. However, multiple factors might better explain why we do not observe an effect of inflammation: 1) disease duration; 2) the effect of dopaminergic medication; 3) life style effects on IL-6; 4) whether IL-6 was determined in serum versus CSF.

First, later disease stages are associated with more neuropsychiatric suffering^65^ and possibly with higher inflammatory tone (although the increase in inflammatory tone might not be linear in PD^66^). Thus, an association with neuropsychiatric suffering and inflammation might be more readily found in later disease stages. Indeed, Lindqvist *et al.*,^44^ who reported increased neuropsychiatric suffering in patients with higher inflammatory concentrations, had an average disease duration that was substantially longer than our sample (i.e. ∼7.5 versus 2 years).

Second, the present study was performed in patients who were ON their normal medication regime, whilst there is evidence that dopaminergic medication alleviates inflammatory side-effects. Felger *et al.*^67^ used levodopa to completely restore striatal dopamine release induced by chronical interferon-α treatment.^67^ Moreover, Bekhbat *et al.*^68^ investigated resting-state functional connectivity between the ventral striatum and vmPFC. They found that only patients with high inflammatory tone (high CRP) showed improvement in resting-state functional connectivity when taking levodopa. In addition, this was associated with a reduction in anhedonia. Thus, levodopa is both able to (partially) restore functional connectivity in the reward circuitry, as well as alleviate neuropsychiatric suffering associated with inflammation^68^, which may potentially explain our negative finding.

Third, although IL-6 is a key cytokine in the inflammatory cascade,^47,48^ and is elevated in prodromal PD,^49^ it might not be optimal as an indicator for inflammatory tone. There are indications that IL-6 is more dependent on state-related changes than disease severity. For example, regular exercise leads to a two and a half fold lower baseline serum concentration for Il-6,^69^ while serum IL-6 can be a 100 fold higher after acute exercise^70^. Regular coffee consumption is associated with 50% higher IL-6 concentrations.^71^ Mindfulness increased the striatal response to rewards, which was associated with reduced IL-6 concentrations.^72^ Hence, these lifestyle factors are likely to have a larger impact on the variance of IL-6 then PD, making it difficult to assess PD related change in inflammatory tone through IL-6.

Last, there might be a divergence between serum and CSF cytokine concentrations. Across fields, multiple studies find dissimilar outcomes for serum and CSF concentrations.^73–75^ Thus, serum-based studies might not readily predict the outcomes of the present work. For example, although Harrison *et al.*^32^ Found a serum IL-6 response after typhoid vaccination, we do not know whether changes in IL-6 concentrations were also reflected in the CSF of these patients, despite it’s ability to modify the RPE. Hence, a correlation between CSF inflammatory response and reward learning might not be found in either samples.

In addition to potential confounding factors that might explain why we did not find a correlation between IL-6 and ICD, there is also substantial between-subject variability in the effects of dopamine of the reward system.^4^ It is possible that such between-subject factors may mask effects for specific subgroups within ICD. For example, the classical inflammation hypothesis indicates decreased reward sensitivity with higher inflammatory tone,^32,76^ due to reduced dopamine availability.^12,23–25^ However, studies found higher inflammatory tone in people with opioid^77^ and alcohol^78^ use disorders, which are disorders that have some resemblance to ICD in PD. This discrepancy, where high inflammatory tone is associated with both reduced reward sensitivity, but higher incidence of addictive behaviour, might be elaborated by future research investigating the relationship between impulsivity and inflammation, where between subject variability in for example type of ICD is considered.

Furthermore, there is still a debate on how inflammation affects reward learning. Inflammation is not consistently associated with altered behavioral outcomes during reward (learning) paradigms. Although there is considerable evidence that inflammation reduces reward sensitivity,^32,76^ there are also multiple contrasting studies who find an increase,^76,79^ or no correlation^80,81^ between reward sensitivity and increased inflammation. This does not indicate that effects of inflammation on the reward system are trivial, but instead points to important inter-individual differences between PD patients.^4^

In the current study, we found a negative correlation between MCP-1 and reward versus punishment learning. MCP-1 is a small cytokine, belonging to the chemokine subfamily (CCL2). It recruits monocytes and memory T-cells and is associated with chronic inflammation in psoriasis, rheumatoid arthritis, MS and obesity.^82^ The finding that higher inflammatory tone is associated with reduced reward versus punishment prediction errors is in line Harrison *et al.,*^32^ who showed reduced reward versus punishment prediction errors after a typhoid vaccination. In addition, MCP-1 has previously been linked with depression in PD.^44^ Patients with major depressive disorder also show reduced reward versus punishment prediction errors.^83,84^ One might thus ask whether reduced reward prediction error signaling mediates the link between inflammation and depression. In fact, in the present work, we found no such correlation between depression and MCP-1, or between depression and reward prediction error signalling.

A methodological issue that we encountered was the inability to measure MIP-1α, IL- 2, IL-12p70, IL-1β, IL-4, IL-17A, IFN-γ and TNF-α. Most likely, the concentrations of these cytokines were too low to be quantified. However we cannot exclude two alternative explanations. First, the samples might have been diluted too much, despite our best efforts to adhere to the protocol. Second, since the protocol has foremost been used on serum samples, it might also require additional optimization for CSF, especially since we used the *high-sensitivity* procartaplex panel.

In summary, we assessed whether reward and punishment learning in PD, as well as depressive and impulsive symptom severity were associated with inflammatory tone in the CSF, particularly IL-6. We found no such effects. When we explored correlations with MCP- 1, we found that higher concentrations of MCP-1 were associated with reduced reward versus punishment learning. However, since this exploratory effect was not corrected for multiple comparisons, it should be replicated in further studies.

## Acknowledgements

We would like to acknowledge the Personalized Parkinson Project team, for their work on data acquisition. In addition, we are very grateful to the kind people who participated in this study.

## Abbreviations

PD: Parkinson’s Disease
ICD: Impulse Control Disorder
PPP: Personalized Parkinson Project
MDS-UPDRS: Movement Disorder Society Unified Parkinson’s disease Rating Scale
RL: Reinforcement Learning
DAT: Dopamine Transporter
RPE: Reward Prediction Error
EV: Expected Value
vmPFC: ventral medial Prefrontal Cortex
BDI: Beck Depression Inventory II
QUIP-rs: Questionnaire for Impulsive-Compulsive Disorders in Parkinson’s disease Rating Scale
MOCA: Montreal Cognitive Assessment
STAI: Anxiety Inventory for Adults
GLM: General Linear Model
ITI: Inter Trial Interval
RLDM: Reinforcement Learning and Decision Making
TFCE: Threshold Free Cluster Enhancement
WM: Working Memory
BH4: Tetrahydrobiopterin
MRI: magnetic resonance imaging
fMRI: Functional magnetic resonance imaging
SDMT: Symbol Digit modalities test
GRO-α: CXCL1
IL: Interleukin
IP-10: Interferon gamma-induced protein 10
MCP-1: monocyte chemoattractant protein 1
MIP-1α: Macrophage Inflammatory Proteins one alfa
MIP-1β: Macrophage Inflammatory Proteins one beta
SDF-1α: stromal cell-derived factor 1 alfa
TNF-α: Tumor necrosis factor alfa
BMI: Body mass index
CSF: Cerebrospinal fluid
BOLD: Blood-Oxygen-Level-Dependent
TR: Repetition Time
ROI: Region Of Interest
PCA: Principal Component Analysis
BRMS: Bayesian Regression Models using ‘Stan’
WSLS: Win-Stay-Lose-Shift
RT: Reaction Time
TFCE: Treshold Free Cluste Enhancement

