## Supplementary material for "Reinforcement learning in Parkinson’s disease is not associated with inflammatory tone"

#### Participants successfully perform the reinforcement learning paradigm

We assessed whether participants showed successful engagement in the instrumental learning task. As expected, accuracy, as measured by choosing the cue with the highest reward probability, increased during the task (main effect of timepoint on $p(cue_{correct})$; brms 95% CI = [0.03 0.05]). Moreover, participants stayed with the same cue more often, if it was rewarded on the previous trial (main effect of outcome on p(stay); brms 95% CI = [0.40 0.63]) an indication of win-stay-lose-shift behavior. Both the increase of accuracy over time and successful engagement in WSLS behavior let us to conclude that participants show adequate task engagement. In addition, we found that overall accuracy was higher in GAIN trials compared to LOSS trials ($p(cue_{correct})$ ~ valence * timepoint); brms 95% CI = [0.003 0.016])).

### Main effects of reward prediction error and expected value

To validate our approach to assess expected value and RPE signals, we assessed the main effect of RPE and EV across all participants. We modeled individual behavior according to a standard Rescorla-Wagner model, to compute both reward prediction error (at time of outcome) and expected value (when the cue is presented). We used a standardized learning rate ($\alpha$=.2) and used the outcomes as parametric regressors in a general linear model [1].

In line with previous work [2], we find highly significant RPE-related brain activity across all subjects and across GAIN and LOSS trials. Both the left (74 voxels, MNI local maximum [-13 7 -9], TFCE = 816.69, $P_{fwe} < 0.001$, supplementary fig. 1A) and right (66 voxels, MNI local maximum [12 4 -9], TFCE = 754.42, $P_{fwe} < 0.001$, supplementary fig. 1A) ventral striatum, as well as a cluster in the medial prefrontal cortex (434 voxels, MNI local maximum [4 49 -2], TFCE = 692.15, $P_{fwe} = 0.001$) show RPE related BOLD response. In addition, we report a cluster in the superior frontal gyrus (60 voxels, MNI local maximum [-16 35 51], TFCE =501.68, $P_{fwe} = 0.002$) and a large cluster in the precuneus and cingulate gyrus (1766 voxels, MNI local maximum [1 -32 33], TFCE = 1082.3, $P_{fwe} < 0.001$). Lastly, we report a valence effect on reward prediction errors; a cluster comprising most of the brain (4945 voxels, MNI local maximum [-24 -98 9], TFCE =11607.33, $P_{fwe} < 0.001$, supplementary fig. 1B), including the ventral striatum, with a clear peak in the visual lobe.

In addition, we replicate a main effect of EV in a ROI analysis of the ventral medial prefrontal cortex (21 voxels, MNI local maximum [1 38 -12], TFCE =70.21, $P_{fwe} = 0.003$, supplementary fig. 1C). The ROI was based on a meta-analysis for EV.[2] The cluster did not reach significance on the whole-brain level. We find no valence effects on EV across all participants.

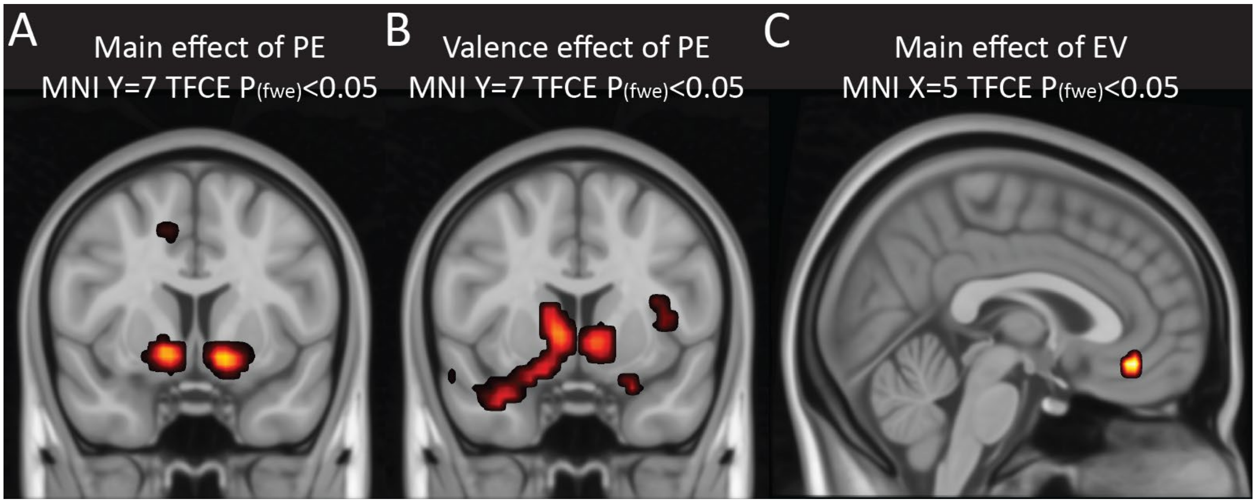


*Supplementary Figure 1 - Main and valence effects of RPE and EV - The variables are calculated by a simple Rescorla-Wagner model and added as parametric regressor to cue (EV) and outcome (PE) onsets. The images were adjusted for family-wise error correction with TFCE p(fwe)<0,05. Reported images are across all participants. A) Whole-brain main effect of reward prediction error B) Whole-brain valence effect (GAIN > LOSS) of reward prediction error C) ROI main effect of EV. ROI was based on a meta-analysis of EV and was comprised of the ventral medial prefrontal cortex [2].*

#### Additional correlations between cytokines and outcome variables

Exploratory, we assessed all possible correlations between the cytokines and clinometric, & RL variables, as presented in Supplementary Table 1. None of these frequentist correlations survive correction for false discovery rate.

We find evidence for a negative correlation between cognitive impairment as measured by the Montreal cognitive assessment and IL-10 (Pearson Rho = -0.34, p=0.029, Bayesian Pearson Rho = -0.33, BF10=3.61, i.e. moderate evidence). IL-10 is an anti-inflammatory marker, indicating increased cognitive decline with decreased reduction of the inflammation response.

Additional classic correlations between cytokines and clinometric factors were not sustained by Bayesian analysis, we found only anecdotal evidence for a correlation between disease duration and IP10 (BF10=1.76), as well as MOCA and SDF-1α (BF10=1.31). The correlation between age and MIP-1β (BF10=0.58), depression (BDI) and RANTES (BF10=0.58) and cognitive decline (MOCA) and the chemokines PCA (BF10=0.56) gave anecdotal evidence for the null hypothesis. Since these factors both do not survive multiple comparison correction and have a BF close to 1, we are stringent to interpret these results.

| | EOTAXIN | GRO-A | IL-8 | IP-10 | MCP-1 | MIP-1B | RANTES | SDF-1A | IL-6 | IL-10 | PCA-CHEMOKINES |
|---|---|---|---|---|---|---|---|---|---|---|---|
| **AGE** | 0.18 | 0.1 | 0.11 | 0.08 | 0.07 | 0.16 | 0.07 | -0.02 | -0.14 | 0.15 | 0.13 |
| **BMI** | 0.06 | -0.09 | -0.16 | -0.02 | -0.01 | -0.12 | 0.03 | 0.03 | 0 | -0.1 | -0.04 |
| **LEDD** | -0.18 | 0.02 | -0.07 | -0.12 | -0.03 | -0.13 | -0.2 | -0.06 | 0.01 | -0.13 | -0.13 |
| **MDS-UPDRS III** | 0.01 | 0.03 | 0.06 | 0.13 | -0.11 | -0.06 | -0.03 | 0.09 | 0 | 0.07 | 0.05 |
| **MONTHS SINCE DIAGNOSIS** | -0.22 | -0.09 | -0.11 | -0.22 | -0.18 | -0.27* | -0.23 | -0.08 | 0 | -0.25 | -0.26* |
| **IMPULSIVITY (QUIP-RS)** | 0.09 | -0.11 | -0.2 | -0.1 | -0.05 | -0.19 | -0.17 | -0.12 | -0.09 | 0.01 | -0.15 |
| **DEPRESSION (BDI-II)** | -0.08 | -0.02 | 0 | -0.01 | -0.15 | -0.05 | -0.27* | 0.1 | 0 | -0.02 | -0.05 |
| **ANXIETY (STAI)** | 0.07 | 0.13 | -0.07 | -0.08 | -0.05 | 0.08 | -0.1 | -0.04 | -0.04 | -0.05 | 0 |
| **COGNITIVE DECLINE (MOCA)** | -0.05 | -0.18 | -0.27* | -0.24* | 0.05 | -0.25* | -0.09 | -0.31* | 0.17 | -0.33* | -0.29* |
| **EV (GAIN + LOSS)** | 0.17 | -0.09 | 0.11 | 0.22 | -0.01 | 0.12 | 0.09 | 0.13 | -0.03 | 0.28 | 0.06 |
| **PE (GAIN + LOSS)** | -0.05 | -0.11 | -0.09 | -0.07 | 0.02 | -0.04 | -0.12 | -0.14 | -0.22 | -0.12 | -0.06 |
| **EV (GAIN > LOSS)** | -0.08 | -0.15 | -0.1 | -0.02 | 0.07 | -0.15 | -0.06 | -0.14 | 0.04 | -0.04 | -0.15 |
| **PE (GAIN > LOSS)** | -0.16 | 0.08 | 0.26* | 0.21 | -0.33** | 0.03 | -0.07 | 0.27* | 0.15 | 0.01 | 0.08 |
| **RESPONSETIME** | [-0.07 0.03] | [-0.04 0.07] | [-0.07 0.04] | [-0.07 0.04] | [-0.02 0.08] | [-0.07 0.04] | [-0.05 0.05] | [-0.07 0.04] | [-0.07 0.04] | [-0.1 0.04] | [-0.06 0.05] |
| **ACCURACY** | [-0.37 0.1] | [-0.23 0.22] | [-0.25 0.22] | [-0.17 0.28] | [-0.29 0.14] | [-0.24 0.21] | [-0.24 0.21] | [-0.22 0.24] | [-0.14 0.32] | [-0.34 0.32] | [-0.24 0.21] |
| **WSLS** | [-0.16 0.06] | [-0.11 0.11] | [-0.08 0.14] | [-0.07 0.15] | [-0.11 0.1] | [-0.09 0.12] | [-0.09 0.14] | [-0.12 0.11] | [-0.12 0.11] | [-0.21 0.14] | [-0.09 0.12] |

*Supplementary Table 2 - Correlation matrix of all possible correlations between the cytokines and the clinometric & RL data.*

### Region Of Interest

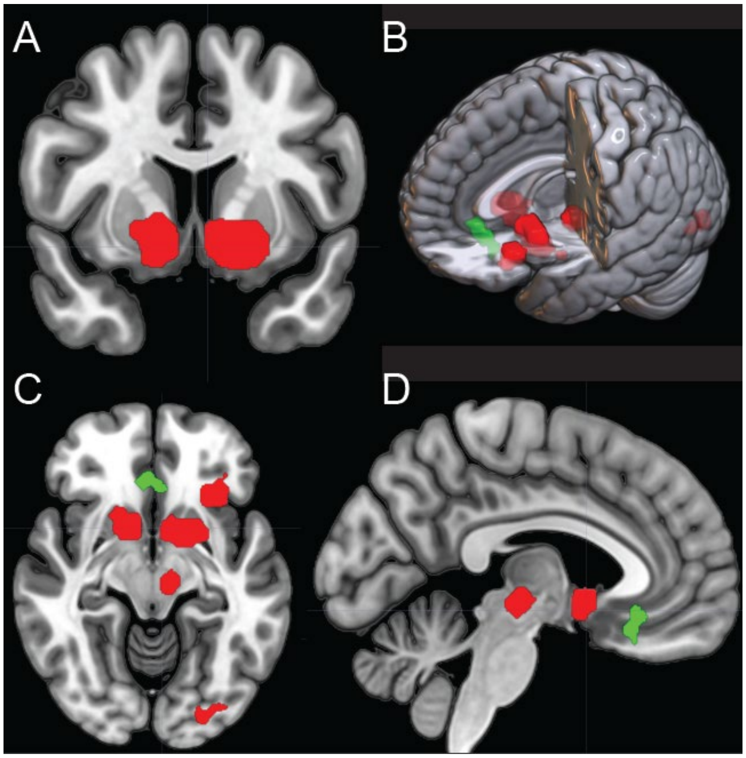


*Supplementary Figure 2 - Masks used for the region of interest analysis - RED: reward prediction error, consisting of the of the striatum, left insula, thalamus, V3 and V4. GREEN: expected value, consisting of a part of the ventromedial prefrontal and orbitofrontal cortex. Masks are based on Chase et al.,* [2] ***(A)*** *MNI X=-5* ***(B)*** *3D render* ***(C)*** *MNI Z=-10* ***(D)*** *MNI Y=8*

### Model space

| | cModelName | cModel | Effect | q25 | q975 | Significant |
|---|---|---|---|---|---|---|
| **1** | clean | correct_response_num ~ valence + (1 + valence \| SubjectNumber) | valence1 | -0.13 | 0.25 | FALSE |
| **2** | ICD_class_IL6 | correct_response_num ~ valence * class_ICD * z_ln_IL_6 + (1 + valence \| SubjectNumber) | valence1:class_ICD1:z_ln_IL_6 | -0.24 | 0.20 | FALSE |
| **3** | ICD_class_chemokines | correct_response_num ~ valence * class_ICD * z_PCA_LN_CHEMOKINES + (1 + valence \| SubjectNumber) | valence1:class_ICD1:z_PCA_LN_CHEMOKINES | -0.29 | 0.21 | FALSE |
| **4** | ICD_IL8 | correct_response_num ~ valence * class_ICD *z_ln_IL_8 + (1 + valence \| SubjectNumber) | valence1:class_ICD1:z_ln_IL_8 | -0.31 | 0.27 | FALSE |
| **5** | ICD_MIP1Beta | correct_response_num ~ valence * class_ICD * z_ln_MIP_1Beta + (1 + valence \| SubjectNumber) | valence1:class_ICD1:z_ln_MIP_1Beta | -0.30 | 0.17 | FALSE |
| **6** | IL6 | correct_response_num ~ valence * z_ln_IL_6 + (1 + valence \| SubjectNumber) | valence1:z_ln_IL_6 | -0.29 | 0.10 | FALSE |
| **7** | Chemokines | correct_response_num ~ valence * z_PCA_LN_CHEMOKINES + (1 + valence \| SubjectNumber) | valence1:z_PCA_LN_CHEMOKINES | -0.27 | 0.11 | FALSE |
| **8** | IL8 | correct_response_num ~ valence *z_ln_IL_8 + (1 + valence \| SubjectNumber) | valence1:z_ln_IL_8 | -0.28 | 0.11 | FALSE |
| **9** | MIP1BETA | correct_response_num ~ valence * z_ln_MIP_1Beta + (1 + valence \| SubjectNumber) | valence1:z_ln_MIP_1Beta | -0.24 | 0.14 | FALSE |
| **10** | Learning_ACC | correct_response_num ~ valence * TimePoint + (1 + valence * TimePoint \| SubjectNumber) | valence1:TimePoint | 0.01 | 0.04 | TRUE |
| **11** | clean | StayShift_num ~ valence * OutcomeM1 + (1 + valence * OutcomeM1 \| SubjectNumber) | valence1:OutcomeM11 | -0.09 | 0.11 | FALSE |
| **12** | ICD_class_IL6 | StayShift_num ~ valence * OutcomeM1 * class_ICD * z_ln_IL_6 + (1 + valence * OutcomeM1 \| SubjectNumber) | valence1:OutcomeM11:class_ICD1:z_ln_IL_6 | -0.11 | 0.14 | FALSE |
| **13** | ICD_class_chemokines | StayShift_num ~ valence * OutcomeM1 * class_ICD * z_PCA_LN_CHEMOKINES + (1 + valence * OutcomeM1 \| SubjectNumber) | valence1:OutcomeM11:class_ICD1:z_PCA_LN_CHEMOKINES | -0.10 | 0.10 | FALSE |
| **14** | ICD_IL8 | StayShift_num ~ valence * OutcomeM1 * class_ICD *z_ln_IL_8 + (1 + valence * OutcomeM1 \| SubjectNumber) | valence1:OutcomeM11:class_ICD1:z_ln_IL_8 | -0.10 | 0.13 | FALSE |
| **15** | ICD_MIP1Beta | StayShift_num ~ valence * OutcomeM1 * class_ICD * z_ln_MIP_1Beta + (1 + valence * OutcomeM1 \| SubjectNumber) | valence1:OutcomeM11:class_ICD1:z_ln_MIP_1Beta | -0.11 | 0.08 | FALSE |
| **16** | IL6 | StayShift_num ~ valence * OutcomeM1 * z_ln_IL_6 + (1 + valence * OutcomeM1 \| SubjectNumber) | valence1:OutcomeM11:z_ln_IL_6 | -0.12 | 0.09 | FALSE |
| **17** | Chemokines | StayShift_num ~ valence * OutcomeM1 * z_PCA_LN_CHEMOKINES + (1 + valence * OutcomeM1 \| SubjectNumber) | valence1:OutcomeM11:z_PCA_LN_CHEMOKINES | -0.13 | 0.05 | FALSE |
| **18** | IL8 | StayShift_num ~ valence * OutcomeM1 *z_ln_IL_8 + (1 + valence * OutcomeM1 \| SubjectNumber) | valence1:OutcomeM11:z_ln_IL_8 | -0.14 | 0.04 | FALSE |
| **19** | MIP1BETA | StayShift_num ~ valence * OutcomeM1 * z_ln_MIP_1Beta + (1 + valence * OutcomeM1 \| SubjectNumber) | valence1:OutcomeM11:z_ln_MIP_1Beta | -0.09 | 0.08 | FALSE |
| **20** | clean | response_time ~ valence + (1 + valence \| SubjectNumber) | valence1 | -0.12 | -0.09 | TRUE |
| **21** | ICD_class_IL6 | response_time ~ valence * class_ICD * z_ln_IL_6 + (1 + valence \| SubjectNumber) | valence1:class_ICD1:z_ln_IL_6 | -0.01 | 0.03 | FALSE |
| **22** | ICD_class_chemokines | response_time ~ valence * class_ICD * z_PCA_LN_CHEMOKINES + (1 + valence \| SubjectNumber) | valence1:class_ICD1:z_PCA_LN_CHEMOKINES | -0.04 | 0.01 | FALSE |
| **23** | ICD_IL8 | response_time ~ valence * class_ICD *z_ln_IL_8 + (1 + valence \| SubjectNumber) | valence1:class_ICD1:z_ln_IL_8 | -0.04 | 0.01 | FALSE |
| **24** | ICD_MIP1Beta | response_time ~ valence * class_ICD * z_ln_MIP_1Beta + (1 + valence \| SubjectNumber) | valence1:class_ICD1:z_ln_MIP_1Beta | -0.03 | 0.01 | FALSE |
| **25** | IL6 | response_time ~ valence * z_ln_IL_6 + (1 + valence \| SubjectNumber) | valence1:z_ln_IL_6 | -0.03 | 0.01 | FALSE |
| **26** | Chemokines | response_time ~ valence * z_PCA_LN_CHEMOKINES + (1 + valence \| SubjectNumber) | valence1:z_PCA_LN_CHEMOKINES | -0.02 | 0.01 | FALSE |
| **27** | IL8 | response_time ~ valence *z_ln_IL_8 + (1 + valence \| SubjectNumber) | valence1:z_ln_IL_8 | -0.02 | 0.02 | FALSE |

| | | | | | | |
|---|---|---|---|---|---|---|
| **28** | MIP1BETA | response_time ~ valence * z_ln_MIP_1Beta + (1 + valence \| SubjectNumber) | valence1:z_ln_MIP_1Beta | -0.02 | 0.01 | FALSE |

*Supplementary Table 3 - Model space*

### Model parameters

*Supplementary Table 4 –* ***Model parameters –*** *An overview of all model parameters used for analysis. Parameter starting with a Z are z-scored.*

| REGRESSOR NAME | REGRESSOR TYPE | REGRESSOR DESCRIPTION | REGRESSOR OPTIONS |
|---|---|---|---|
| **VALENCE** | Binary | Trial type of the current variable | GAIN (1) or LOSS (0) |
| **GROUP (I.E. MEDICATION)** | Binary | Group to which the subject belonged. For each option, a separate model was run. | ON (1) vs OFF (0)<br>ON (1) vs HC (0)<br>OFF (1) vs HC) (0) |
| **OUTCOMEM1** | Binary | Indicating whether the previous trial was either rewarded (+€10 for GAIN trials or €0 for LOSS trials) or punished (€0 for GAIN trials or €-10 for LOSS trials). | Rewarded (1) or punished (0) |
| **Z_BDI** | Continuous | Summary score of the BDI questionnaire, indicating the degree of depression | |
| **Z_QUIPRS_TOT** | Continuous | Summary score of the QUIP-rs questionnaire, indicating the degree of impulsivity | |
| **Z_STAI** | Continuous | Summary score of the STAI questionnaire, indicating the degree of anxiety in the current state (opposed to trait) | |
| **CLASS_QUIPRS** | Binary | Classification of clinical ICD based on (CITE) | ICD (1) and non-ICD (0) |
| **CLASS_BDI_SIMPLIFIED** | Binary | Classification of clinical depression | Depressed (1) and non-depressed (0) |
| **CORRECT_RESPONSE_NUM** | Binary | Shows whether the correct cue was chosen on a given trial | Correct (1) or incorrect (0) |
| **STAYSHIFT_NUM** | Binary | Shows whether the same cue as the previous trial of that trial type was chosen for a given trial | Same (1) or other (0) |
| **RESPONSE_TIME** | Continuous | Reaction time between cue presentation and response in milliseconds | |
| **Z_LEDD** | Continuous | Levodopa Equivalent Daily Dose | |
| **USEAGO** | Binary | Whether or not this participant actively uses dopamine agonists | Yes (1) or No (0) |

| | | |
|---|---|---|
| **SYMBOLDIGIT** | Continuous | Composed score of the Symbol and Digits task, indicating general cognitive capability |
| **MONTHSSINCEDIAG** | Continuous | Number of months between the measurement and diagnosis |
